# Stabilized Supralinear Network: Model of Layer 2/3 of the Primary Visual Cortex

**DOI:** 10.1101/2020.12.30.424892

**Authors:** Dina Obeid, Kenneth D. Miller

## Abstract

Electrophysiological recording in the primary visual cortex (V1) of mammals have revealed a number of complex interactions between the center and surround. Understanding the underlying circuit mechanisms is crucial to understanding fundamental brain computations. In this paper we address the following phenomena that have been observed in V1 of animals with orientation maps: 1) surround suppression that is accompanied by a decrease in the excitatory and inhibitory currents that the cell receives as the stimulus size increases beyond the cell’s summation field; 2) surround tuning to the center orientation, in which the strongest suppression arises when the surround orientation matches that of the center stimulus; and 3) feature-specific suppression, in which a surround stimulus of a given orientation specifically suppresses that orientation’s component of the response to a center plaid stimulus. We show that a stabilized supralinear network that has biologically plausible connectivity and synaptic efficacies that depend on cortical distance and orientation difference between neurons can consistently reproduce all the above phenomena. We explain the mechanism behind each result, and argue that feature-specific suppression and surround tuning to the center orientation are independent phenomena. Specifically, if we remove some aspects of the connectivity from the model it will still produce feature-specific suppression but not surround tuning to the center orientation. We also show that in the model the activity decay time constant is similar to the cortical activity decay time constant reported in mouse V1. Our model indicates that if the surround activates neurons that fall within the reach of the horizontal projections in V1, the above mentioned phenomena can be generated by V1 alone without the need of cortico-cortical feedback. Finally, we show that these results hold both in networks with rate-based units and with conductance-based spiking units. This demonstrates that the stabilized supra-linear network mechanism can be achieved in the more biological context of spiking networks.

## Introduction

Electrophysiological recording from cells in the primary visual cortex (V1) reveal that visual stimuli presented outside the classical receptive field (CRF) of a neuron (the surround) can modulate the neuron’s response to a stimulus present in its CRF (the center) in complex ways. The degree and direction of modulation depends on the distance between the center and surround, the contrasts of the stimuli, their relative orientations, etc. (Akasaki et al., 2002; Bair et al., 2003; Cavanaugh et al., 2002; Sceniak et al., 1999; Shen et al., 2007; Sillito et al., 1995; Wang et al., 2009). Which of these modulations are carried by V1 lateral connections, and which require top-down signals from higher visual areas is still largely unknown. Understanding the underlying circuit mechanisms is crucial to understanding fundamental brain computations.

To address these mechanisms, we build a spatially-extended, biologically-constrained model of layer 2/3 of V1 of animals with orientation maps. We investigate whether a set of key phenomena that have been reported in V1 can be consistently generated by lateral connections alone, without the need of cortico-cortical feedback. We find that lateral connections are sufficient provided that specific conditions for connectivity and synaptic efficacies are met. Therefore, our model makes testable predictions about the structure of the underlying circuit.

We first address surround suppression. We show that our model can successfully reproduce surround suppression in similar strength and with similar contrast dependence to that observed in layer 2/3 of V1 of animals with orientation maps. Furthermore, this suppression is accompanied by a decrease in both excitatory and inhibitory conductances that the cell receives, as reported in Ozeki et al. (2009). In order to achieve this, in addition to structured connectivity, the network must locally have strong connections. Although many studies have shown that surround suppression in V1 can be mediated through lateral connections (Rubin et al., 2015), this is the first demonstration of the accompanying decrease in received inhibition as well as excitation in a spatially extended (spatially two-dimensional) model.

We then investigate two new phenomena: (1) The strongest suppression arises when the surround orientation matches that of the center stimulus, even when the center orientation is not optimal for the cell (Shushruth et al., 2012; Trott and Born, 2015) and (2) A surround with orientation matching the orientation of one component of a plaid center stimulus more strongly suppresses the response of the matching component (Trott and Born, 2015). We show that phenomena (1) and (2) can be generated within V1 if the surround falls with the reach of V1 lateral connections. We find that to match (1), local connectivity, in addition to being strong, must be broadly tuned for orientation; however to match (2) this additional requirement is not needed. This leads us to conclude that effects (1) and (2) are independent.

We further show that in the model the activity decay time constant is fast, similar to the cortical activity decay time constant reported by Reinhold et al. (2015). Finally, we show that our results hold in networks with conductance-based spiking units as well as rate units. This demonstrates that the Stabilized-Supralinear Network mechanism described in Ahmadian et al. (2013) and Rubin et al. (2015) can arise in the more biological context of a spiking neural network (see also Sanzeni et al. (2020a,b)).

## Model

### Model Overview

To investigate the computational role of V1 lateral connections, we build a 2-dimensional spatially extended model of layer 2/3 of the primary visual cortex of animals with orientation maps. Retinotopic position changes smoothly across both spatial dimensions, while preferred orientation of neurons is determined by their position in the orientation map. The Cortical Magnification Factor (CMF), which expresses how many mm of cortex represents one degree in visual angle, constrains the size of a neuron’s receptive field (RF), as we describe below.

The connectivity in the model is broadly constrained by biological data. Neurons in V1 layer 2/3 are found to form dense axonal projections at distances of a few hundred *μm*, and sparse long range horizontal projections that target cells of similar orientation preferences. These long range connections, which can reach up to 3 *mm* in cat and 10 *mm* in monkey, arise from excitatory cells, and give rise to the patchy connectivity observed in V1 (Amir et al., 1993; Bosking et al., 1997; Stettler et al., 2002). In comparison, inhibitory cells primarily form short range connections.

We first present results from a rate-based model. The units in the rate-based model are taken to have an expansive or supralinear, power-law transfer function (Albrecht and Hamilton, 1982; Albrecht, 1991; Carandini et al., 1997, 1999; Finn et al., 2007; Hansel and Van Vreeswijk, 2002; Heeger, 1992; Miller and Troyer, 2002), as expected for neurons whose spiking is driven by input fluctuations rather than by the mean input (Hansel and Van Vreeswijk, 2002; Miller and Troyer, 2002). Rubin et al. (2015) and Ahmadian et al. (2013) showed that when neural-like units have such a power-law transfer function, responses with nonlinear behaviors observed in visual cortex emerge due to network dynamics. The authors called this mechanism the Stabilized Supralinear Network (SSN). They showed that the SSN mechanism can explain normalization and surround suppression and their nonlinear dependencies on stimulus contrast, which are observed across multiple sensory cortical areas.

To verify that our results are robust and independent of the neuron model, we also build a conductance-based spiking neural network model, and show that all our key results still hold. This shows as well that the SSN mechanism can be realized with spiking neurons.

### Model Details

We use a grid of 75 × 75 grid points. We place one excitatory cell (E), and one inhibitory cell (I) at each location on the lattice, and thus have 5625 E cells and 5625 I cells in the network. Even though we use a 50/50 E/I ratio, we believe that our main results will not change if we take the E/I ratio to be 80/20. In unpublished work we studied SSN behavior in spiking networks consisting of 1152 E cells and 288 I cells (E/I ratio of 80/20), and found that the network behavior was consistent with SSN predictions in the parameter regime we studied (strongly suppressive regime, *i.e*. with Ω_*E*_ < Ω_*I*_ < 0, using parameters defined in Ahmadian et al. (2013)). We take the map to represent 16×16 degrees of visual space, with position in visual space varying linearly across the map, and assume a Cortical Magnification Factor (CMF) of 0.5 mm/deg. Thus the grid represents 8.0 × 8.0 mm of cortex, with each grid interval representing 0.213 degrees and 107 μm of cortical distance. We use periodic boundary conditions; our results are independent of that condition. This is verified by removing periodic boundary conditions, and adjusting the weight efficacy matrix to compensate for the lost connections.

We superpose on the grid an orientation map, specifying the preferred orientations of cells at the corresponding grid points (Fig. 1A). The orientation map is generated randomly using the method described in Kaschube et al. (2010) (their supplementary materials, Eq. 20). To summarize, we superpose *n* complex plane waves to form a function *z*(x) of two-dimensional spatial position x:

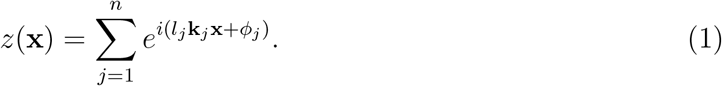

Here, k_*j*_ = *k*(cos(*jπ*/*n*), sin(*jπ*/*n*)), with signs *l_j_* ∈ {+1, −1} and phases *ϕ_j_* ∈ [0,2*π*) randomly chosen. Writing *z*(x) = *r*(x)*e*^*i*Φ(x)^ for real amplitude *r*(x) and phase Φ(x), we take the preferred orientation at each grid point x to be Φ(x)/2. We use a map spatial frequency of 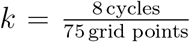, *i.e*. a map with on average 8 full periods of the orientation map across the length or width of the grid, and *n* = 30. The orientation map is not periodic, so there is a discontinuity in orientation at the grid borders, although the retinotopy and intracortical connections wrap around. In our results, we report on cells sampled away from the boundary (20 < *x* < 60,20 < *y* < 60, in terms of the grid coordinates that go from 1 to 75 in each dimension) to avoid boundary effects.

The excitatory cells form long range connections, while the inhibitory cells form short range connections. The connection strength from a unit of type *Y* and grid position *b* to a unit of type *X* at position *a, X, Y* ∈ {*E,I*}, is written 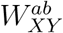. Let the units at *a* and b have positions x_a_ and x_b_, respectively, and preferred orientations *θ_a_* and *θ_b_*. The connection strength is given by 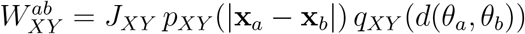, where *d*(*θ_a_,θ_b_*) is the shortest angular distance around a 180° circle between the two orientations. Here, *p_XY_*(x) describes the dependence of strength on the spatial distance between the units (measured as the shortest distance across the grid with periodic boundary conditions), while *q_XY_*(θ) describes the dependence on the difference between their preferred orientations measured as shortest distance around the circle of orientations. The function *p_XY_*(x) is specified as follows: for projections of excitatory cells, *p_XE_*(x) is 1 for distances *x* ≤ *L_o_*, and then decays as a Gaussian with standard deviation *σ_XE_*. *L_o_* = 324 μm, *σ_EE_* = 324 μm and = 642 μm. For projections of inhibitory cells, *p_XI_*(x) is Gaussian with standard deviation *σ_EI_* = *σ_II_* = 216 μm. For all cells regardless of pre- or postsynaptic type, the function *q_XY_*(*θ*) has the form of a Gaussian with a non-zero baseline: 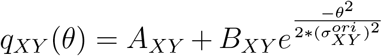. For projections of I cells and of E cells at distances less than *L_o_, A_XI_* = *A_XE_* = 0.2, *B_XI_* = *B_XE_* = 0.8 and 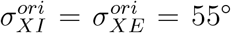. For projections of excitatory cells at distances greater than *L_o_, A_XE_* = 0.14, *B_XE_* = 0.86 and 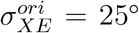. The constants *J_XY_* are, for I projections, *J_EI_* = 0.0528 and *J_II_* = 0.0288; for E projections, at distances less than *L_o_, J_EE_* = 0.072 and *J_IE_* = 0.06, while at distances greater than *L_o_, J_EE_* = *J_IE_* = 0.036. We point out that the heterogeneity in the network comes from the underlying orientation map.

**Figure 1:**
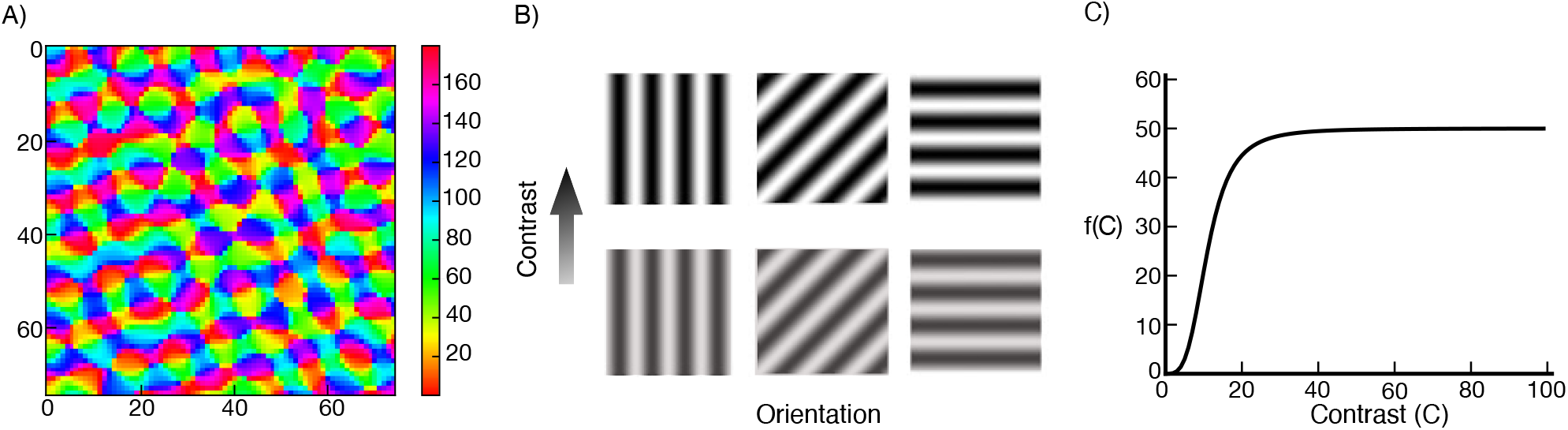
(A) Orientation map, the color corresponds to the cells preferred orientation. (B) Gratings with different orientations and contrasts. (C) External input as a function of the stimulus contrast (Eq. 3).

We choose the connectivity parameters so that the connectivity profile agrees with experimental findings. We choose the *J_XY_* such that 1) the network is in a strong sublinear regime (see Result 1 for more details) and 2) with increasing stimulus size, the loss of excitatory input to inhibitory cells from nearby surround-suppressed excitatory cells is greater than their gain in excitatory input from far away excitatory cells, which is necessary for the net inhibition received by excitatory cells to decrease with surround suppression. We constrain the rest of the model parameters by experimental data to make the model more biologically plausible.

We ignore stimulus features like spatial frequency and phase, and consider only three features: contrast, orientation and size (Fig. 1B). The cells in the model behave like ideal complex cells, in that their response to a drifting grating is static in time. Spatially, each cell has a circularly symmetric Gaussian receptive field with standard deviation *σ_rf_* = 0.09°. The external input to a neuron located at position (*x_o_,y_o_*) with preferred orientation *θ_o_*, from a stimulus of contrast C and orientation *θ_s_* that is centered at (*x_s_,y_s_*) and is uniform for a diameter of *ℓ* degrees about the center (and zero contrast outside), is given by

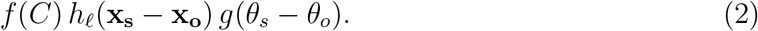

Here *f*(*C*) is a Naka-Rushton function given by

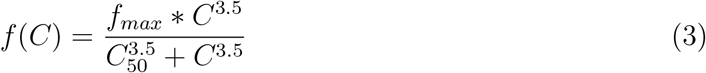

with *f_max_* = 50 and *C_50_* = 11 (Fig. 1C). *h_ℓ_*(**x**) is the integral of the product of the Gaussian classical receptive field with a sharp edge stimulus. It is given by:

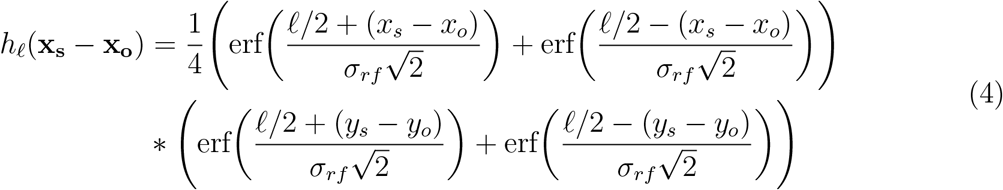

where erf(x) is the error function defined as 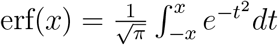. The function *g* is defined by

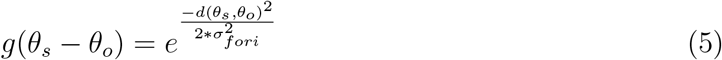

with *σ_fori_* = 20°. To define the equations for the rate model, we let 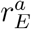 be the rate of the excitatory neuron at position *a*, and 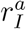 similarly. Both receive the same external input 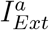. The rate equations are:

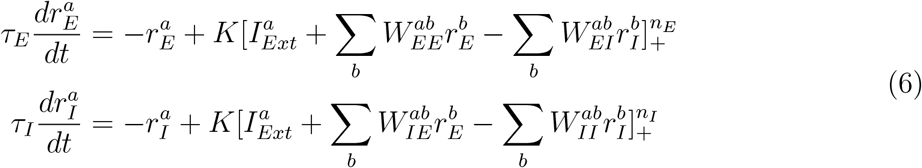

where [*x*]_+_ = max(0,*x*). The excitatory cells’ time constant *τ_E_* = 10*ms*, and the inhibitory cells’ time constant *τ_I_* = 6.67*ms*. We use, *K* = 0.01, *n_E_* = *n_I_* = 2.2. 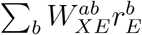 is the recurrent excitatory input to neuron *X^a^* where *X* = {*E,I*}. Similarly, 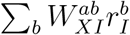 is the recurrent inhibitory input.

For the conductance-based model, the equations of motion of the membrane potential and the conductances for each cell are identical for E and I cells. For a cell at *a* of type *X*, the equations are (we omit specifying the type *X* for the dynamical variables and parameters that don’t differ between the two types):

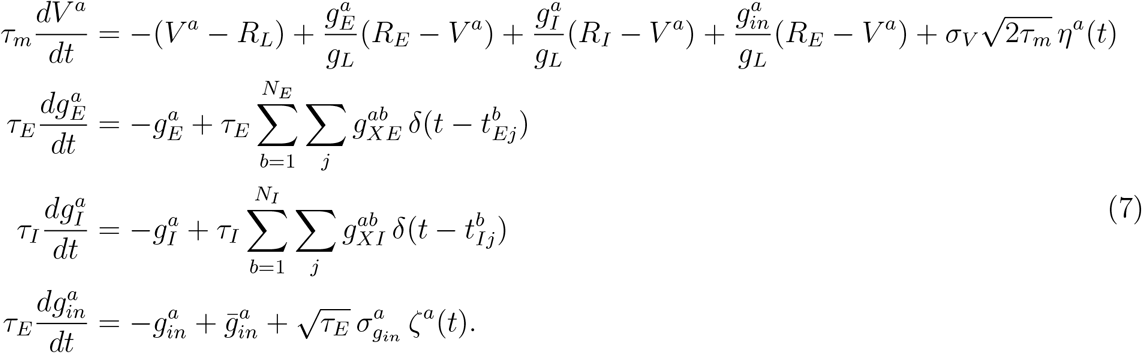

Here *V^a^* is the membrane potential of the given cell at a. *τ_m_* is the membrane potential time constant. 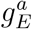 is the excitatory AMPA-like conductance, 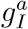 is the inhibitory GABA-like conductance and 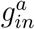 is the excitatory input conductance from outside the network. *R_E_*, *R_I_*, and *R_L_* are reversal potentials of the excitatory, inhibitory, and leak conductances. *τ_E_* and *τ_I_* are the time constants of the excitatory and inhibitory conductances. 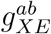 is the conductance of the synapse of the excitatory cell at *b* to the given cell at *a*, similarly 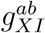 is the conductance from the inhibitory cell at *b*, and 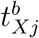 is the time of the *j^th^* spike of the cell of type *X* at *b*. *δ*(*x*) is the Dirac delta function. Each cell in the network receives input from *N_input_* external spiking cells, where *N_input_* is a large number. We assume the spike trains are Poisson and invoke the Central Limit Theorem to approximate the input to the cell 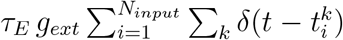 by a stochastic process with mean 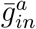 and variance 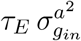, where *ζ^a^* is white Gaussian noise with < *ζ^a^*(*t*)*ζ^a^*(*t′*) >= *δ*(*t − t′*). The stochastic dynamics will lead 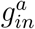 to have a mean 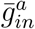 and a variance 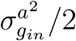 (Tuckwell, 1988), where

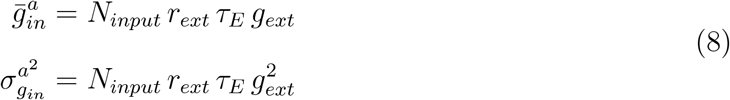

where *g_ext_* is the amplitude of the conductance evoked when a single external cell spikes, and *r_ext_* is the firing rate of the external cells given by Eq. 2. We assume the membrane potential is noisy, and model the noise as white Gaussian noise. *η^a^*(*t*) is a Gaussian random variable with mean 0 and variance 1, and *σ_V_* is the standard deviation of the membrane potential fluctuations. In the simulations we set *σ_V_* = 6.85 *mV* to get spontaneous activity similar to what has been reported in (Chen et al., 2009; Gur and Snodderly, 2008; Ringach et al., 2002). In the model the mean spontaneous activity of the excitatory cells is about 1.5 Hz, and of the inhibitory cells is about 3 Hz. The parameters for both E and I cells are as follows: *τ_m_* = 15 *ms*; *τ_E_* = *τ_I_* = 3*ms*; *R_L_* = −70 *mV*; *R_E_* = 0 *mV*; *R_I_* = −80 *mV*; *g_L_* = 10 *nS*; *N_input_* = 200 and *g_ext_* = 0.1 *nS*. We take threshold voltage *V_th_* = −50 *mV* and after-spike rest voltage to be 6 *mV* below threshold, *V_r_* = −56 *mV*, as in Troyer and Miller (1997). After the cell spikes, it goes into a refractory period with *τ_re_f* = 3 *ms*.

Similar to the rate model, the conductance values are given by 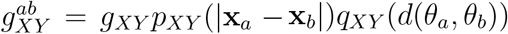. The parameters *g_XY_* are: *g_EI_* = 3.3*nS* and *g_II_* = 2*nS*; at distances less than *L_o_, g_EE_* = 1.8 *nS* and *g_IE_* = 1.76 *nS*; and at distances greater than *L_o_, g_EE_* = 0.7 *nS* and *g_IE_* = 0.65 *nS*. Again *L_o_* = 324 μm as in the rate model.

To measure the strength of the surround suppression in the network, we compute the suppression index (SI) defined as:

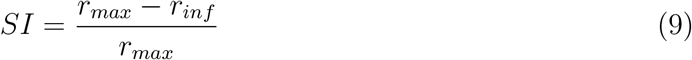

where *r_max_* is the response to a stimulus size that elicits maximum response, and *r_inf_* is the response for a very large stimulus. To measure whether the presence of a surround stimulus facilitates or suppresses the response of a cell compared to its response to a center-only stimulus, we define a modified suppression index:

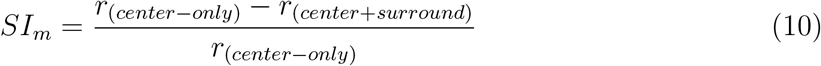

*SI_m_* negative means facilitation, while *SI_m_* positive means suppression.

In experiments on the surround tuning to the center orientation, we fix the center stimulus diameter, and the inner and outer annulus diameters to (1.3°, 4.3°, 21.6°) and set both stimuli contrasts to 100. In these experiments we record the activity of a single neuron as we vary the stimulus orientations, and we roughly pick the largest annulus inner diameter at which the phenomena is still observed. This corresponds to an annulus inner radius of 2.15° or 1.1 *mm* which is roughly the span of E-to-I monosynaptic connections. In feature-specific suppression experiments we fix the center stimulus diameter, and the inner and outer annulus diameters to (1.7°, 3.9°, 21.6°). In these experiments we follow the procedure in Trott and Born (2015) to make our results directly comparable with experimental data. Thus, we use a slightly bigger center stimulus to obtain a better fit of the population rates (see Result 3 for more details). The contrast is set to 16.4 (representing 80% of the maximal input strength), for each component of the plaid as well as the surround in the rate model, and to 50 (representing 99.5% of the maximal input) in the conductance-based model.

In all experiments, cells are sampled randomly from locations away from the boundary. We first randomly pick 100 locations within the region we define as away from the boundary (20 < *x* < 60, 20 < *y* < 60). Cells in all experiments are randomly picked from those 100 locations.

## Results

We first check that our network is functioning as an SSN, by checking for several salient SSN behaviors. The SSN shows a transition, with increasing input strength, from a weakly coupled, largely feedforward driven regime for weak external input, to a strongly coupled, recurrently-dominated regime for stronger external input (Ahmadian et al., 2013). This transition can account for many aspects of summation of responses to two stimuli and of center-surround interactions and their dependencies on stimulus contrast (Rubin et al., 2015). Our network shows the characteristic signs of this transition (Rubin et al., 2015): the net input a neuron receives grows linearly or supralinearly as a function of external input for weak external input, but sublinearly for stronger external input (Fig. 2A); this net input is dominantly external input for weak external input, but network-driven input for stronger external input (Fig. 2B); the network input becomes increasingly inhibitory with increasing external drive (Fig. 2C); and a surround stimulus of a fixed contrast can be facilitating for a weak center stimulus, but becomes suppressive for stronger external drive to the center (Fig. 2D).

**Figure 2:**
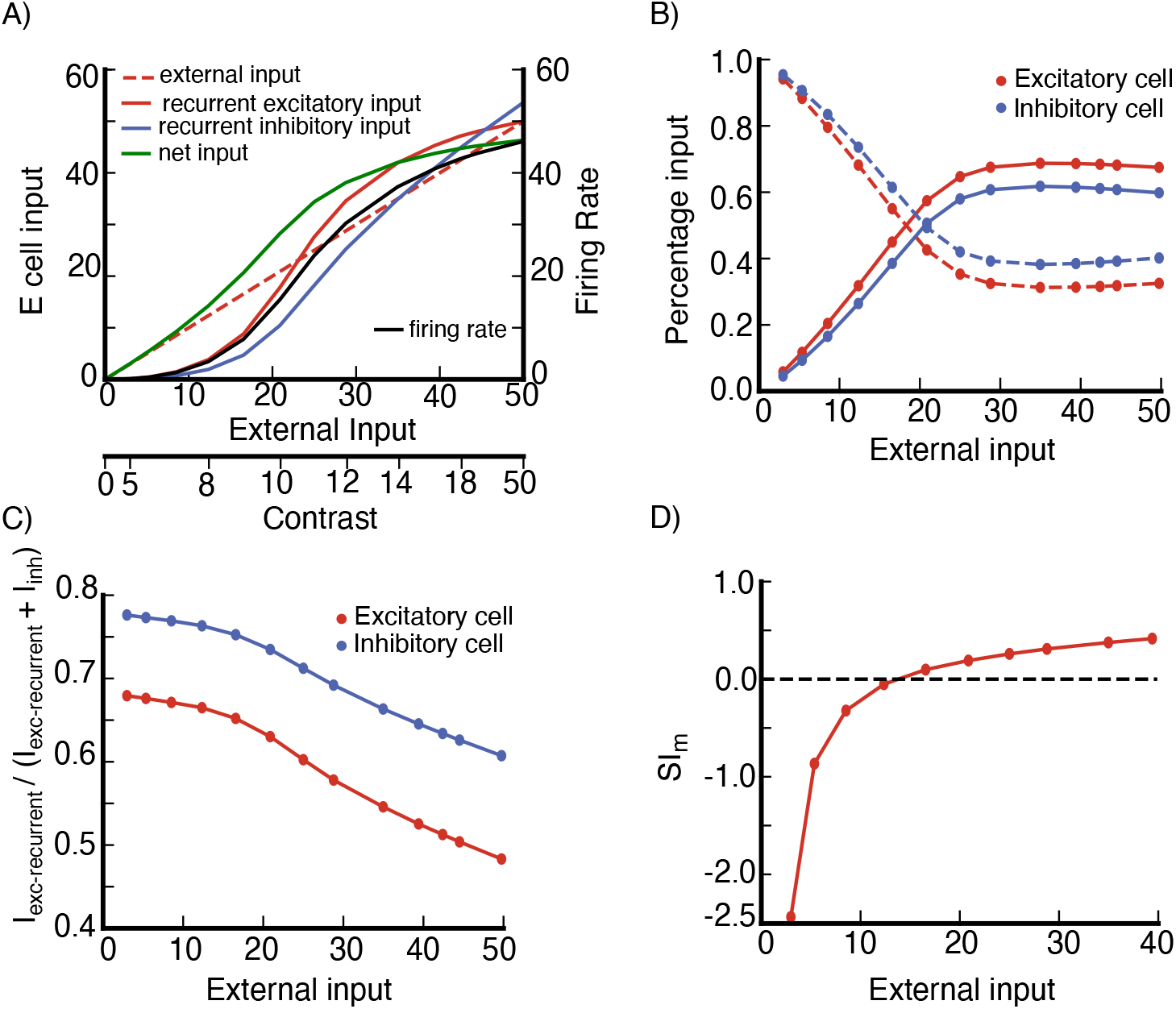
Stabilized Supralinear Network (SSN) behavior of the model network. (A) Inputs to an excitatory (E) cell and its firing rate vs external input, the cell is at a randomly selected grid location (see section Model Details). Stimulus contrast level corresponding to external input is shown on the bottom axis. The net input is defined as (*I_ext_* + *I_exc–recurrent_ − I_inh_*), where *I_ext_* is the external input to the cell, *I_exc–recurrent_* is the cell’s recurrent excitatory input from the network, and *I_inh_* its recurrent inhibitory input from the network, *I_inh_* is defined to be positive, see section Model Details for the expressions of *I_exc–recurrent_* and *I_inh_*. (B) Percentage of external and network inputs as a function of external input for the excitatory cell in (A) and an inhibitory cell at the same grid location (dashed line is external input, solid line is network input). Here, the total input is defined as (*I_ext_* + *I_exc–recurrent_* + *I_inh_*), and the network input is (*I_exc–recurrent_* + *I_inh_*). (C) *I_exc–recurrent_*/(*I_exc–recurrent_* + *I_inh_*) as a function of external input for the excitatory and inhibitory cells in (A,B). In panels (A-C) we use a stimulus of diameter 2.16° centered on the cell’s retinotopic position and with the cell’s preferred orientation. (D) Surround Facilitation to Suppression transition: a near surround can be facilitating or suppressing depending on the center stimulus contrast. *SI_m_* negative means facilitation, while *SI_m_* positive means suppression (see section Model Details, Eq. 10 for the definition of *SI_m_*). In panel (D) the data is from an excitatory cell at a randomly selected grid location(see section Model Details); surround stimulus has contrast *C* = 12, and inner and outer diameters 0.865° and 4.32° respectively; the center stimulus diameter is 0.65°; both center and surround stimuli are centered on the cell’s retinotopic position, with the cell’s preferred orientation.

We then explore whether lateral connections in V1 are capable of generating several phenomena that emerge due to center-surround interaction.

### Result 1: Surround Suppression

We first investigate surround suppression, a widely studied phenomena in V1 and other sensory areas in multiple species (Angelucci et al., 2017). Ozeki et al. (2009) showed in anesthetized cat V1 that, after presenting an optimal center-only stimulus, presentation of an iso-oriented surround stimulus decreased firing rates, and decreased both the inhibition and excitation neurons receive. Adesnik (2017) similarly showed that inhibition as well as excitation were decreased by surround suppression in awake mice. Ozeki et al. (2009) showed that suppression of inhibition as well as excitation required that the network be an *inhibition stabilized network*, or ISN, meaning that, if inhibition were frozen and could not respond dynamically, the excitatory subnetwork would be unstable. Rubin et al. (2015) demonstrated a circuit model with one spatial dimension in which surround suppression was accompanied by a decrease of inhibition as well as excitation, and showed contrast dependence like that seen in visual cortex. They also studied a model with two spatial dimensions that was in a different parameter regime but that showed similar behaviors. However, since Rubin et al. (2015) was published, we discovered that the 2-D spatial model of V1 studied there did not show a decrease in inhibition received with surround suppression, and to our knowledge no other 2-D spatial model of V1 has shown this. We investigate the conditions under which surround suppression can emerge in a 2-d spatially extended model of layer 2/3 of V1 with a decrease in inhibition received. More specifically, what structure of connectivity and synaptic efficacies can achieve this? Before addressing this question, we first show that our model replicates surround suppression behavior and its contrast dependence observed in Rubin et al. (2015).

To study surround suppression, we record the firing rates of a cell in the network, as we vary the diameter of a high contrast stimulus centered on the cell’s retinotopic position and with orientation identical to the recorded cell’s preferred orientation. In the model both excitatory (E) and inhibitory (I) cells are surround-suppressed. However, excitatory cells are more strongly surround suppressed than inhibitory cells, as illustrated by an E and I cell at a randomly selected grid location (see section Model Details) (Fig. 3A,B) and by the average size tuning across 80 E and 80 I cells (Fig. 3C) for a high contrast stimulus, C=16.4. Accordingly, the summation field sizes – the size of a stimulus driving optimal response, before further increase in size causes response suppression – of E cells are smaller than those for I cells (Fig. 3D).

**Figure 3:**
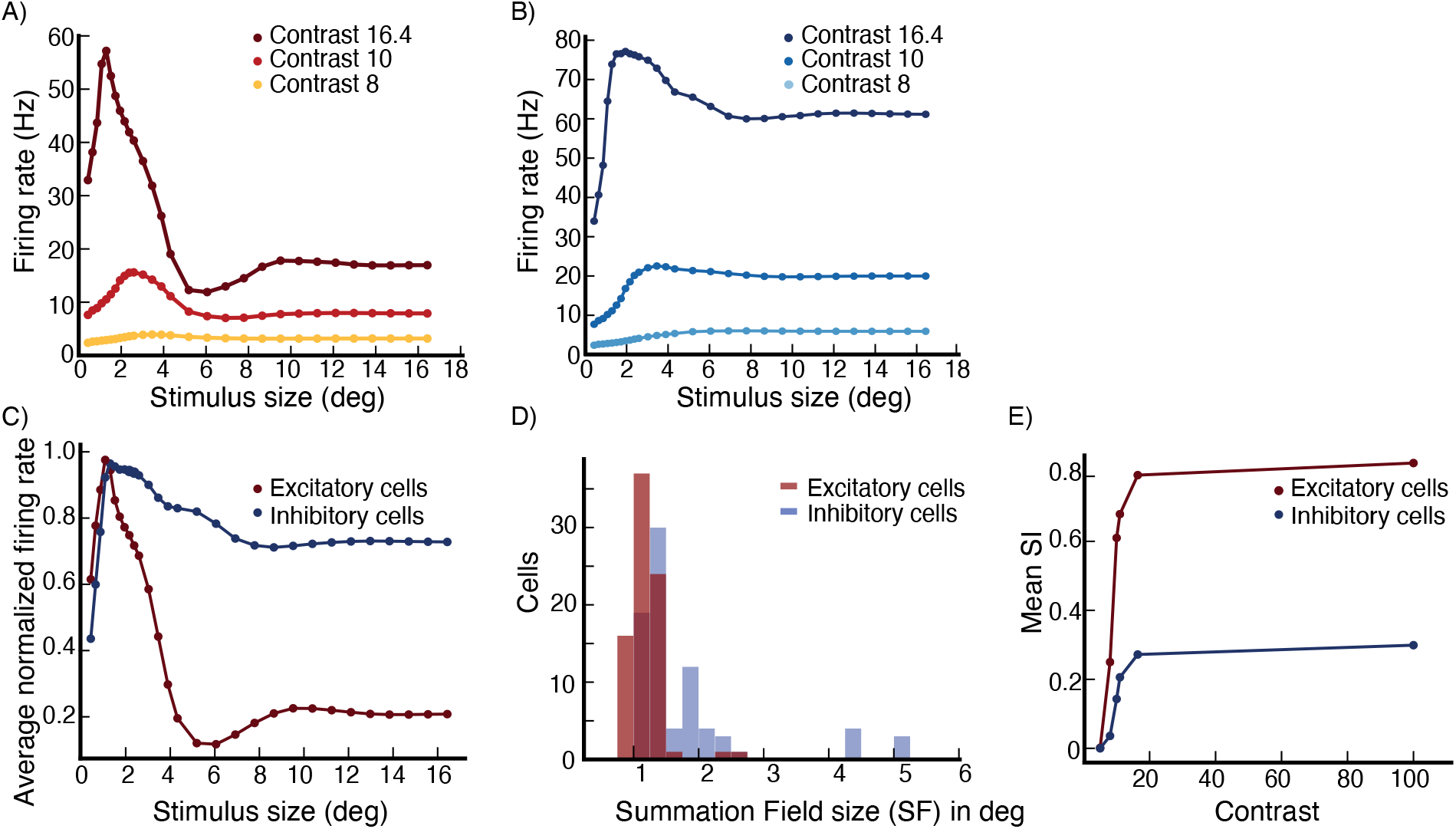
Surround suppression. (A,B) The firing rates of an excitatory (E) cell (A) and an inhibitory (I) cell at the same grid location (B) vs. stimulus size. Different colors correspond to different stimulus contrast levels, high (C=16.4; external input 80% of maximal), medium (C=10; external input 42% of maximal) and low (C=8; external input 25% of maximal). The cells are at a randomly selected grid location (see section Model Details). (C) The average firing rate of 80 E cells at randomly selected grid locations (see section Model Details), and of 80 I cells at the same grid locations, after normalizing each cell’s rates so that its peak rate is 1.0, vs. stimulus size for a high contrast stimulus (C=16.4). (D) The distribution of Summation Field sizes (SFS) of the E and I cells used to produce panel (C), the mean SFS for the E cells is 1.14deg and for the I cells is 1.75 deg. (E) The mean suppression index of the E cells and I cells used to produce panel (C) versus stimulus contrast, the mean Suppression Index (SI) for E and I cells changes from little or no suppression (low SI’s) for very weak stimuli, to stronger suppression (higher SI’s) for stronger stimuli, with E cells showing much stronger suppression than I cells. The error bars are too small to show properly, they are of order 10^−2^ or smaller.

We repeat the above experiment with different contrast levels. The strength of surround suppression increases with increasing stimulus contrast (Fig. 3A,B). The mean suppression index (SI) increases from little or no suppression for weak contrasts to stronger suppression for stronger contrasts (Fig. 3E). For a relatively high contrast stimulus (C=16.4, representing 80% of the maximal input strength), the mean suppression index (SI) is 0.79 for the E cells and 0.27 for the I cells (where 0 is no suppression and 1 is complete suppression). Similarly, the summation field size shrinks with increasing contrast, as we illustrate for E cells in Fig. 4A,B, as in (Sceniak et al., 1999). The summation field sizes of I cells behave similarly.

**Figure 4:**
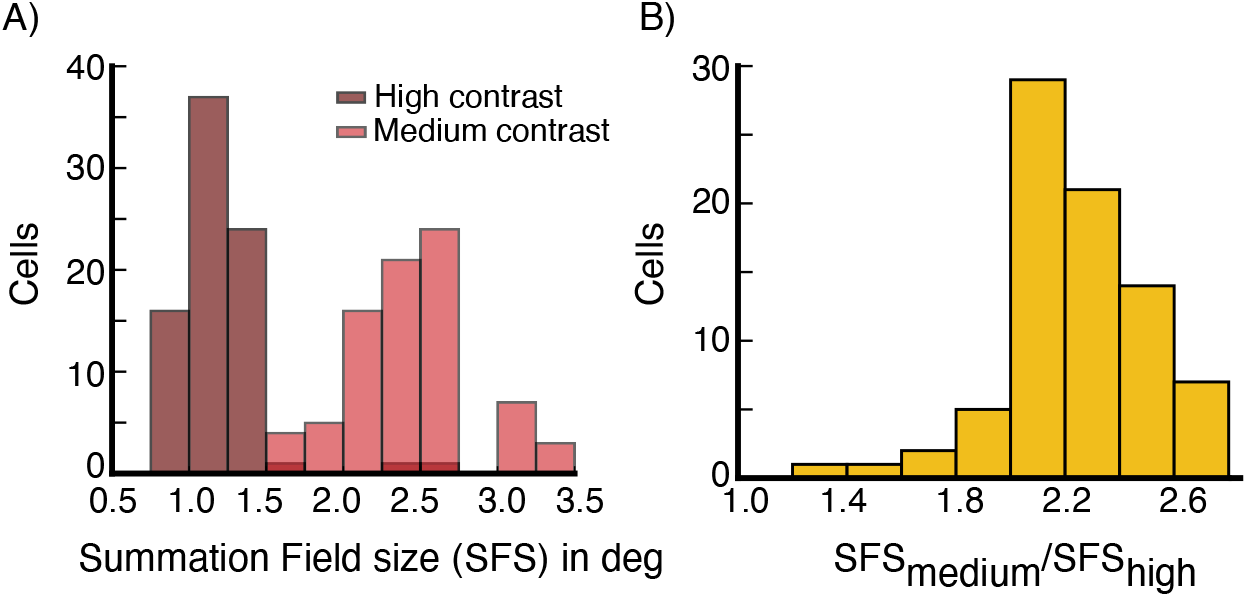
Surround suppression, summation field sizes. (A) The distribution of summation field sizes for 80 excitatory (E) cells (same E cells used to produce panel Fig. 3C), at contrast C=16.4 (dark red color) and contrast C=10 (light red color). (B) The distribution of the ratio of the summation field sizes in (A). The summation field size of all cells is smaller for the higher contrast stimulus.

We examine whether surround suppression in the network is accompanied by a decrease in excitation and inhibition, as reported by Ozeki et al. (2009), rather than simply being due to ramping up of inhibitory input. The size tuning of the excitatory and inhibitory input currents to the E cell in Fig. 3A at high contrast (C=16.4) reveals that both currents indeed show surround suppression (Fig. 5A). We then look at the average size tuning of these currents across cells, after normalizing each cell’s curve for each current to have a peak of 1. Both E cells (Fig. 5B) and I cells (Fig. 5C) show surround suppression of both their excitatory and their inhibitory currents.

**Figure 5:**
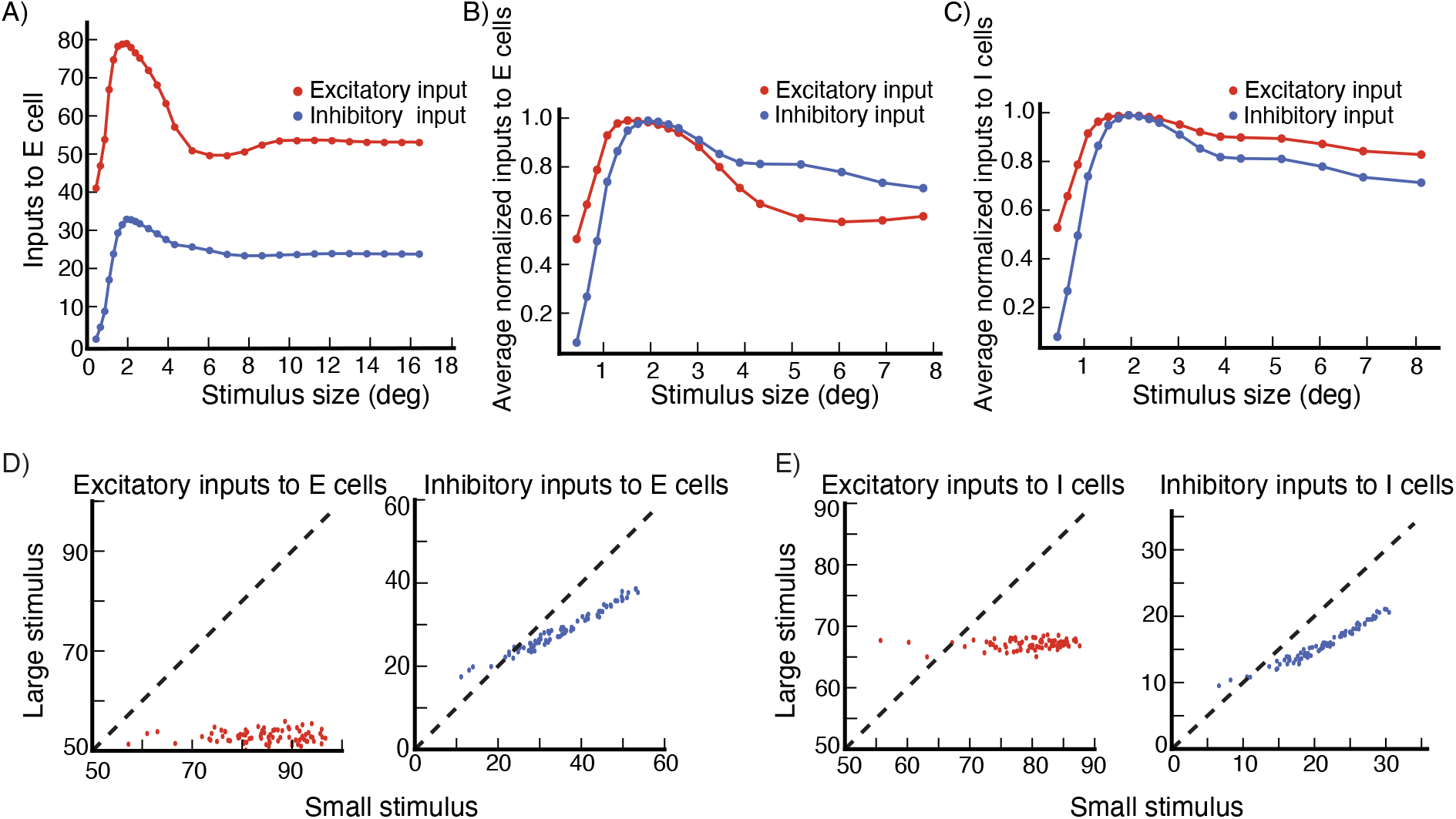
Surround suppression, inputs to cells. (A) The excitatory (red) and inhibitory (blue) total input to the excitatory (E) cell in Fig. 3A, shown for the high contrast stimulus (C=16.4; external input 80% of maximal), both show surround suppression. (B,C) The size tuning of the averaged normalized excitatory and inhibitory inputs (each normalized to have peak value 1) to excitatory (E) cells (B) and inhibitory (I) cells (C) for contrast C=16.4 (same cells used to produce panel Fig. 3C). Note the change in horizontal axis between panels (A) and (B,C). (D) The excitatory and inhibitory inputs to E cells (same E cells used to produce panel Fig. 3C) for a large stimulus (for which all the cells are surround suppressed) are shown vs. their values for a small stimulus (with size given by the average size that yields maximal response across cells). Panel (E) is the same as (D) but for I cells (same I cells used to produce panel Fig. 3C). Stimulus contrast C=16.4.

We then wish to directly compare, across cells, the currents for a small, nearly-optimally-sized stimulus to those for a large, suppressive stimulus. To compare to experiments, there is now a problem to be solved: as modelers we know the exact stimulus size that gives peak response, and can compare inhibition received for that size to inhibition received for a large size. However experimenters do not know the optimal size, and must choose some size in that vicinity, which may evoke less inhibition than the peak (see Fig. 3). Thus, if we choose the optimal size for comparison, we may bias our results towards seeing a decrease in inhibition, compared to experimental procedures. To avoid this, we follow a procedure similar to that of Ozeki et al. (2009). We measure the excitatory and inhibitory inputs for a small stimulus size with diameter *d_s_* around which the cells respond close to maximally, and for a very large stimulus at which all cells are surround suppressed. We take *d_s_* to be equal to the median of all stimulus diameters for which the sampled cells respond maximally. The results are entirely similar if *d_s_* is taken to be the mean rather than the median.

Using this procedure, for excitatory inputs and for inhibitory inputs to E and to I cells, we plot the input current at small stimulus size vs. the current at large stimulus size (Fig. 5D,E). In Fig. 5D we plot the excitatory inputs and inhibitory inputs respectively to 80 excitatory cells, at small stimulus size against those at large stimulus size. Both excitatory and inhibitory inputs are smaller for the large suppressive stimulus. Fig. 5E shows the same data for 80 inhibitory cells. Thus, for both excitatory and inhibitory cells in the model, surround suppression is accompanied by a decease in excitation and inhibition that the cell receives.

While we cannot exhaustively search all parameters, in our explorations of parameters, we have found the surround suppression of inhibitory as well as excitatory input to depend on two elements of the connectivity. First, locally, roughly over distances of about *L_o_* (the distance over which lateral connections are most dense, see section Model Details), the cells must be strongly enough connected so that, as the stimulus size increases, the local circuit around the recorded cell goes through the SSN transition from being mainly driven by the feedforward input to being dominated by recurrent currents. This occurs through the increase in effective synaptic weights with increased external drive to the network due to the expansive, supralinear neuronal input/output function, which is the fundamental mechanism underlying the SSN (see (Ahmadian et al., 2013; Rubin et al., 2015) for a detailed description of the SSN mechanism). At the transition, the growth of effective excitatory synaptic strengths is sufficient that the excitatory subnetwork becomes unstable by itself (Ahmadian et al., 2013), but the network is stabilized by feedback inhibition. This means that the local circuit becomes an inhibition stabilized network (ISN), which was the condition for surround suppression of inhibitory input identified in (Ozeki et al., 2009), based on the ISN mechanism initially identified by (Tsodyks et al., 1997). We put the network into a particular regime of the SSN that is thought to be most strongly non-linear, though we don’t know if this is necessary since we did not do an exhaustive parameter search. This regime, using parameters defined in Ahmadian et al. (2013), is defined by Ω_*E*_ < 0 (in particular, we use Ω_E_ < 0 < Ω_*I*_) where, for equal inputs to the excitatory and inhibitory cells as we use here, Ω_*E*_ = *W_II_* − *W_EI_* and Ω_*I*_ = *W_IE_* − *W_EE_*, with *W_XY_* the total synaptic weight from units of type Y to a unit of type X. This produces a regime in which responses saturate with increasing external input (and ultimately would supersaturate for sufficiently strong external input). In particular, for our connectivity parameters, Ω_*E*_ = −0.49 ± 0.01 and Ω_*I*_ = 3.59 ± 0.03.

The second element we have found critical is that the ratio of projection strength of long-range horizontal connections to I cells vs. E cells must increase with increasing distance, that is, the E-to-I connections must be effectively longer range than E-to-E connections. Furthermore, the excitatory input received by I cells from far away E cells should not be large compared to the excitatory input they receive from nearby excitatory cells. Then, with increasing stimulus size, the loss of excitatory input to I cells from surround suppression of nearby E cells can exceed the gain of excitatory input from far away E cells, causing the I cells to be surround suppressed. Note that, in our model (as in (Rubin et al., 2015)), the I cells have larger summation fields than the E cells (Fig. 3C,D). This means that there is an intermediate range of stimulus sizes for which inhibitory firing rates continue on average to increase with stimulus size, while excitatory cells are surround suppressed. With further increase in stimulus size, both E and I cells are suppressed.

### Result 2: Surround Tuning to the Center Orientation

Cells in V1 are found to be suppressed maximally when the surround stimulus orientation matches the orientation of the center stimulus, regardless of whether that orientation matches the cell’s preferred orientation (Shushruth et al., 2012; Sillito et al., 1995; Trott and Born, 2015). This might enable the cell to detect orientation discontinuities or help in foreground-background separation. A similar behavior has been observed for other stimulus features, such as spatial frequency and velocity (Shen et al., 2007).

A previous model (Shushruth et al., 2012) showed that this behavior could arise if cells received strong, weakly tuned excitatory and inhibitory input from the local network, while the surround drove more strongly tuned inhibition of the excitatory cells and excitation of the inhibitory cells. Then, when the center stimulus differed from the recorded cell’s preferred orientation, the cell would receive a great deal of local recurrent excitation and inhibition from cells preferring the stimulus orientation, which would be the most strongly activated cells. A surround stimulus matched to the center stimulus would most strongly target these most activated cells. Withdrawal of input from those cells would then cause greater suppression of the firing of the recorded cell than would direct suppression from a surround stimulus at the cell’s preferred orientation. Therefore, suppression would be greatest when the surround stimulus orientation matched the center stimulus orientation.

We use similar reasoning here, but now in the context of the SSN model with power-law rather than linear-rectified input/output functions. Our long-range projections are excitatory onto both E and I cells, whereas in (Shushruth et al., 2012) they were inhibitory onto E cells and excitatory onto I cells. In addition, the model of (Shushruth et al., 2012) was not recurrent, because the input from one cell to another was simply determined by the difference in their preferred orientations, regardless of the firing rate of the presynaptic cell (and thus was a constant, independent of the stimulus); and the surround input to a cell was determined only by the difference between the cell’s preferred orientation and the surround stimulus orientation, and not by the firing rates of lateral cells responding to the surround stimulus. We use a recurrent model.

We record the firing rate of cells in the network for different center orientations. For each center orientation, we then present a stimulus in the surround, rotate its orientation, and record the cell’s firing rate for each center-surround orientation configuration. In an excitatory cell (Fig. 6), we study tuning to center orientation absent a surround (black curves) and then tuning to surround orientation for a fixed center stimulus (red curves). The most suppressive surround orientation (minimum of red curve) is pulled strongly toward the center orientation (red asterisk) as the center orientation is varied from −20° relative to preferred orientation (Fig. 6A), to preferred (Fig. 6B), to +20° relative to preferred (Fig. 6C). Similar results are seen more generally in 67 excitatory cells at randomly selected grid locations (Fig. 7). The surround orientation producing maximum suppression is pulled strongly towards the center orientation (Fig. 7A,C-D) and in most cases is within 10° of the center orientation (Fig. 7B).

**Figure 6:**
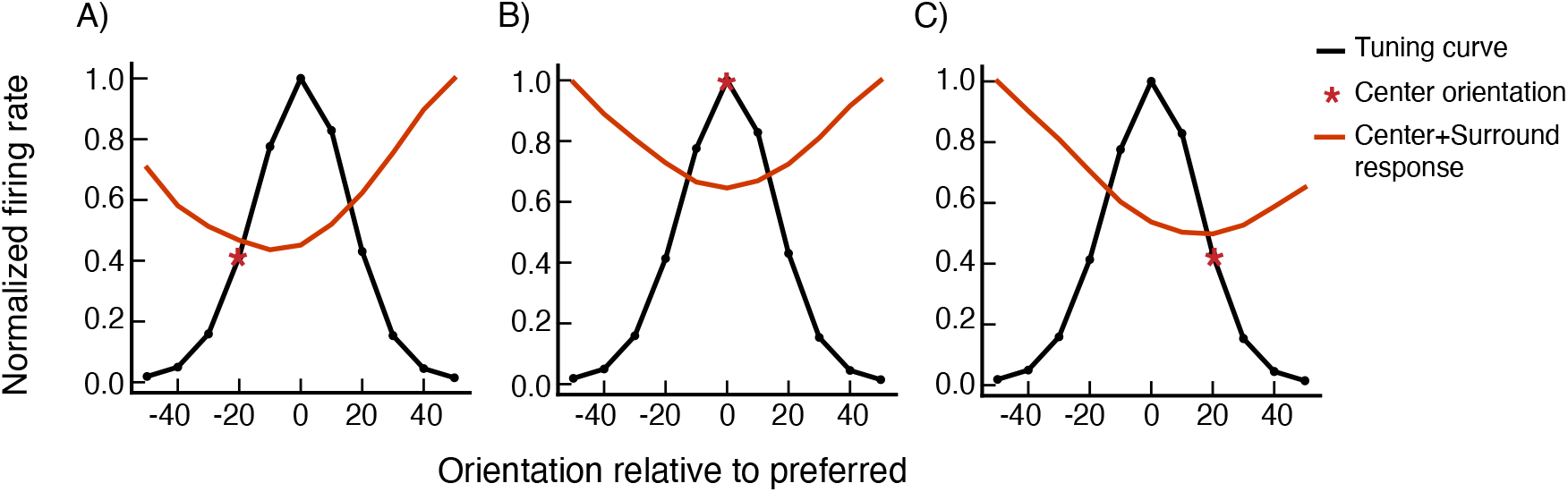
Surround tuning to the center orientation in an excitatory (E) cell. The orientations are relative to the cell’s preferred orientation. The black curves show the cell’s orientation tuning curve for a center-only stimulus (i.e., firing rate vs. center orientation), normalized so the maximum response is 1.0. The red curves show the similarly normalized tuning to surround orientation for a fixed center stimulus. In each panel, the red asterisk marks the fixed center orientation: (A) center at preferred minus 20°, (B) center at preferred and (C) center at preferred plus 20°.

**Figure 7:**
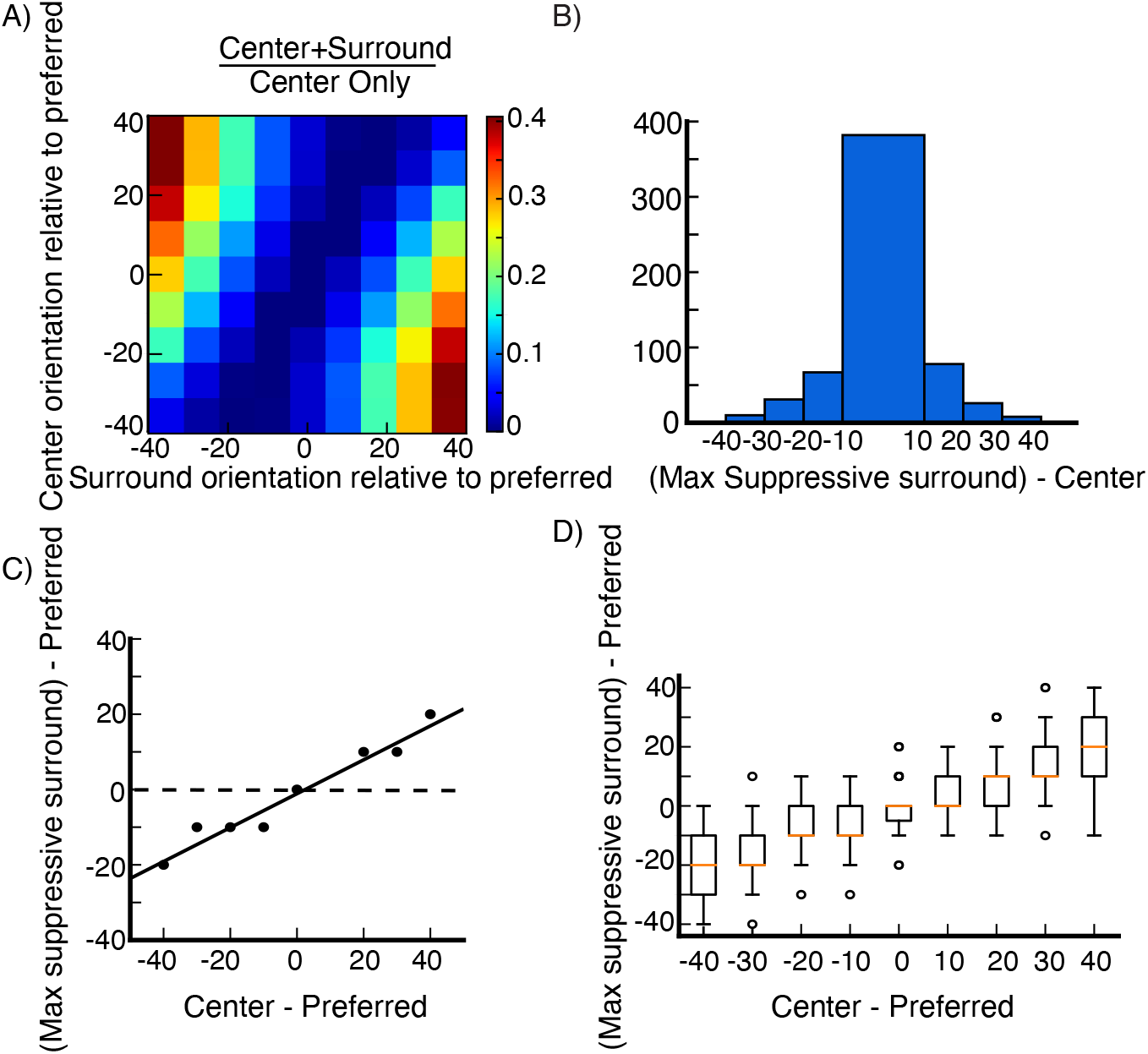
Surround tuning to the center orientation. Surround tuning in 67 excitatory (E) cells at randomly selected grid locations (see section Model Details). Both center and surround orientations are varied from preferred minus 40° to preferred plus 40° in increments of 10°. (A) Average surround modulation map. For each cell, the map is obtained by dividing center-surround responses by the corresponding center-only responses, each row has its minimum value subtracted. (B) Histogram of the difference between the orientation of the surround that maximally suppresses the cell’s firing rate, and the center stimulus orientation. The data is pooled over all cells and center orientations. (C) Orientation of the surround that maximally suppresses the cell’s firing rate plotted against the center stimulus orientation, averaged over all cells. (D) Whisker plot of orientation of the surround that maximally suppresses the cell’s firing rate against the center stimulus orientation, the box extends from quantile Q1 to Q3, the orange line is the median. The upper whisker extends to last datum less than Q3 + k*IQR, similarly, the lower whisker extends to the first datum greater than Q1 - k*IQR, where IQR is the interquartile range (Q3-Q1) and k=1.5, the circles represent the outlier data. In (A), (C), (D), orientations are shown relative to preferred.

As described above, surround tuning to the center orientation arises due to the strong, broadly tuned local connectivity profile in orientation space, along with the more sharply tuned surround input, which causes maximal input to the cell to come from cells preferring the center stimulus rather than from cells with the same preferred orientation as the recorded cell. This makes suppression targeted to cells preferring the center stimulus more potent than suppression targeted to cells preferring the recorded cell’s own preferred orientation.

### Result 3: Feature-Specific Surround Suppression

Surround suppression in V1 is not blind to the center stimulus, as we have just seen. This also manifests in the *feature-specificity* of surround suppression (Trott and Born, 2015): if multiple stimuli are present in the cell’s center, the surround more strongly suppresses the response component driven by the center stimulus whose orientation matches the surround’s. We test whether V1 lateral connections can mediate such computation. We follow the procedure described in Trott and Born (2015).

We record the firing rate of a small population of neurons to each of two oriented gratings. For each stimulus, we fit the average response vs. preferred orientation across the population with a Von Mises function, call these functions P_1_ and P_2_ (Fig. 8A). We then record the population’s firing rates to a center plaid stimulus, the superposition of the two individual gratings. If the two gratings differ by, *e.g*., 60°, we will call this a 60° plaid or a plaid angle of 60°. We fit the population’s response to the plaid stimulus as a linear combination of the two components, R_plaid_= w_1_ P_1_ + w_2_ P_2_.

**Figure 8:**
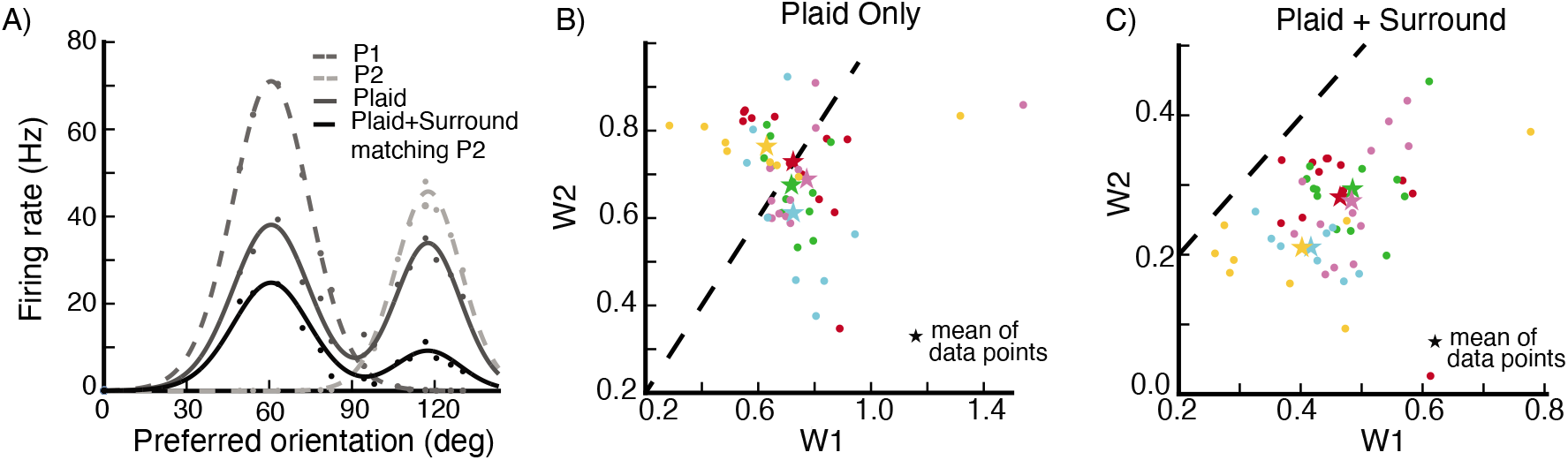
Feature-specific surround suppression. (A) The firing rate of a small population of neurons in response to a center stimulus. The neurons are binned in 5° bins according to their preferred orientation. The dots are the data points, and the lines are the Von-Mises-function fits to the data. The medium gray points and dashed line are the population response to the first component of the plaid (P_1_). The light gray points and dashed line are the population response to the second component of the plaid (P_2_). The dark gray points and solid line are the population response to the plaid. The black points and solid line are the population response to the plaid in the presence of a surround stimulus whose orientation matches the plaid’s second component. (B,C) values of w_1_ and w_2_ (the weightings in fitting the plaid population response to a weighted sum of the two component responses), for a 60° center plaid stimulus, shown for 12 different plaid rotations (every 10°), recorded from five different populations (indicated by colors). The populations are centered around randomly selected grid locations (see section Model Details). Missing data points imply that we can not find a good fit of the data for certain stimulus configurations. The star symbols are the mean values of w_1_ and w_2_ for each location. (B) Responses to plaid center stimulus only. (C) Responses to plaid center stimulus in the presence of a surround stimulus with orientation equal to the plaid’s second component. Dashed lines are unit diagonals, along which w_1_=w_2_

We then introduce a surround stimulus whose orientation matches the second component of the plaid center stimulus, and measure the new values of w_1_ and w_2_. We repeat the same procedure as we rotate the plaid, each time matching the surround stimulus to the orientation of the 2nd plaid component. For responses to a 60° plaid alone, w_1_ and w_2_ on average have about equal strength, but the addition of a surround stimulus matched to the second plaid component suppresses w_2_ much more than w_1_ (Fig. 8B,C).

We carry out this experiment for different plaids, with plaid angles [−60°, −30°, 0°, 30°, 60°, 90°]. The mean values of w_1_ and w_2_ across all of these plaids cluster around w_1_=w_2_ for the plaid stimulus alone (Fig. 9A), but are heavily shifted towards w_1_ when the surround stimulus matched to the second plaid component is added (Fig. 9B). In Fig. 9C we show the mean values of data points in Fig. 9A,B for each plaid angle. In the absence of a surround stimulus there is no difference between w_1_ and w_2_. When a surround stimulus with orientation matching the plaid’s second component is introduced, both components of the plaid are suppressed, however, we clearly see that the second component is suppressed more. Hence, the surround stimulus suppress mostly the center stimulus component that has similar feature. In our model, this phenomena emerges because long range lateral connections connect cells with similar preferred orientations. The surround stimulus mostly excite cells with preferred orientation close to its orientation. These cells in turn will mostly suppress the cells at the center which have similar preferred orientation. Finally, we point out that the results remain the same if we repeat the same experiment and record from a single cell rather than from a small local population.

**Figure 9:**
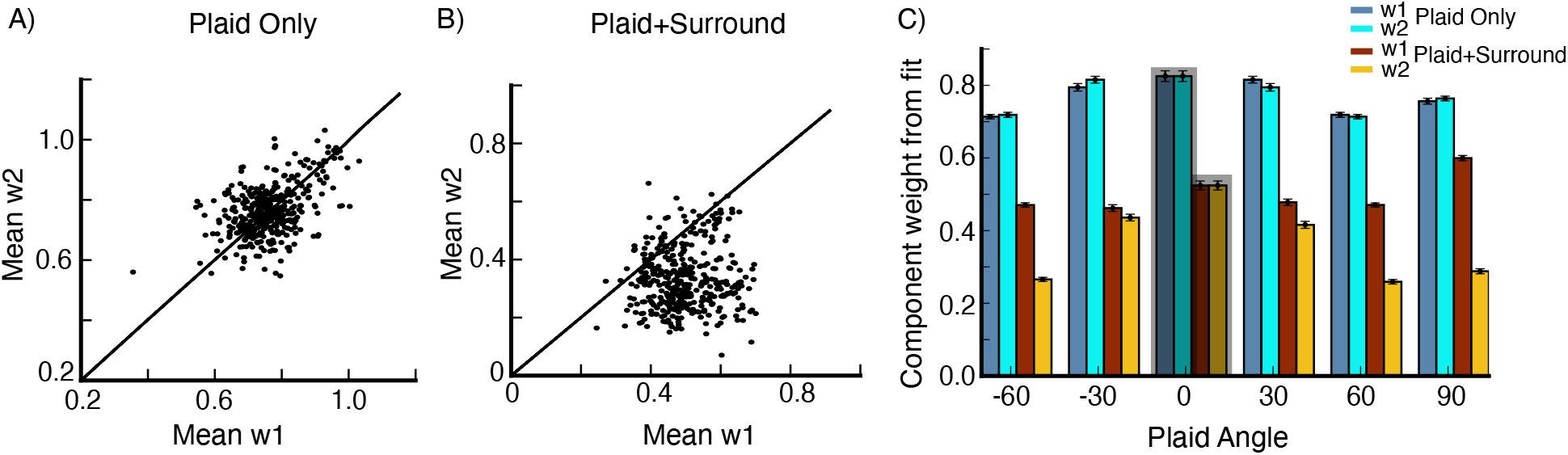
Feature-specific surround suppression. (A,B) Mean w_1_ is plotted against mean w_2_ for plaid angles [−60°, −30°, 30°, 60°, 90°] from 80 populations, centered around 80 randomly selected grid locations (see section Model Details). The mean values of w_1_ and w_2_ are obtained from averaging data for different rotations of the plaid (A) for plaid center stimulus only (B) for plaid center stimulus in the presence of surround stimulus with orientation equal the plaid’s second component. (C) Mean values of the data points in (A) and (B) for different plaid angles, we also include the data for plaid angle 0°, error bars are the s.e.m.

### Result 4: Activity Decay Time

While exciting V1 with a visual stimulus, Reinhold et al. (2015) abruptly silenced the thalamic input to V1, by silencing the lateral geniculate nucleus (LGN) through optogenetic stimulation of the thalamic reticular nucleus (TRN). They showed that, after thalamic silencing, the cortical activity in V1 exhibited a fast decay, two orders of magnitude faster than the decay after visual stimulus offset. The authors called this decay time after thalamic silencing the cortical decay function (CDF). The CDF across all V1 layers was of the order of 10 ms, in particular for multiunits the CDF + s.e.m. was L2/3: 9.8 ± 1.7 ms, L4: 9.0 ± 2.2 ms, L5a: 8.9 ± 1.3 ms, L5b: 15.7 ± 2.5 ms and L6: 7.6 ± 1.5 ms. The CDF was almost the same in awake and anesthetized mice. Furthermore, the authors found that the CDF was independent of the stimulus contrast.

We test if the dynamics in our network are in agreement with what has been reported. Since we only model layer 2/3 in V1, silencing LGN is equivalent to the removal of the feedforward input in our model. We record the activity of a cell for two stimulus sizes 2° and 10°, each at two contrast levels, high contrast (C=17) and low contrast (C=9). In all cases, the feedforward input is removed at 200 ms. To obtain the activity decay time constant, we fit the decaying activity with an exponential function. We find the decay time constant for the excitatory cells to be roughly 10 ms, which is in agreement to what has been reported in Reinhold et al. (2015) as we show in Fig. 10. For 50 excitatory cells at randomly selected grid locations, the activity decay time constant (mean ± s.e.m.) for a 2° stimulus at high contrast is 10.04 ± 0.01ms and at low contrast is 9.99 ± 0.02 ms, and for a 10° stimulus at high contrast 9.75 ms and at low contrast 9.76 ms (when we do not report it, the s.e.m. is too small). The activity decay time constant is almost independent of the stimulus contrast level and size. It is roughly given by the excitatory cells’ time constant in the model. Similarly, the activity decay time scale for the inhibitory cells is roughly given by the inhibitory cells’ time constant. For 2° stimulus at high contrast the inhibitory cells activity decay time constant is 6.93 ± 0.01ms and at low contrast 6.73 ± 0.04 ms, and for the 10° stimulus at high contrast it is 6.69 ± 0.03 ms and at low contrast 6.98 ± 0.05 ms.

**Figure 10:**
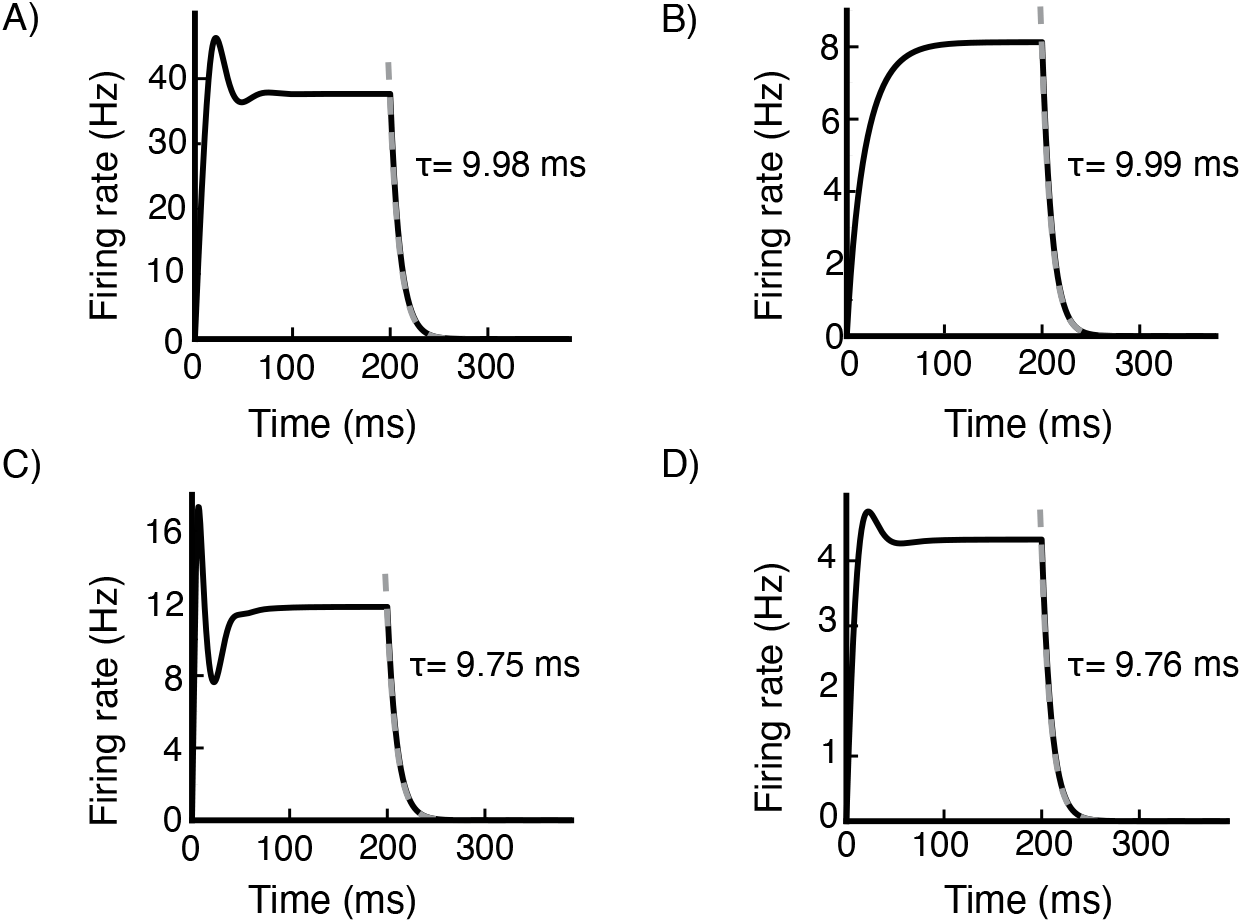
Activity decay time. The time response of an excitatory (E) cell at a randomly selected grid location (see section Mode Details) for various stimulus conditions. (A) and (B) 2° size stimulus at high contrast (C=17) and low contrast (C=9) respectively. (C) and (D) 10° size stimulus at high contrast (C=17) and low contrast (C=9) respectively. The feedforward input is removed at 200 ms. The activity decay time constant is obtained by fitting an exponential function to the decaying activity. The activity decay time constant is roughly independent of the stimulus contrast level and size.

### Result 5: Conductance-based Spiking Model

To test whether our results depend on the neuron model, we replace the rate units in the network with conductance based spiking units (Eq. 7 in section Model Details). To make the model more biologically realistic, we assume the excitatory and the inhibitory cells have spontaneous activity levels of 1.5 Hz and 3 Hz respectively. We first show how the input currents to a cell and its firing rate change with external drive (Fig. 11A,B). The net input current a neuron receives increases rapidly as a function of external current for weak external input, but sublinearly for stronger external input (Fig. 11A). The network input becomes increasingly inhibitory with increasing external drive (Fig. 11C).

**Figure 11:**
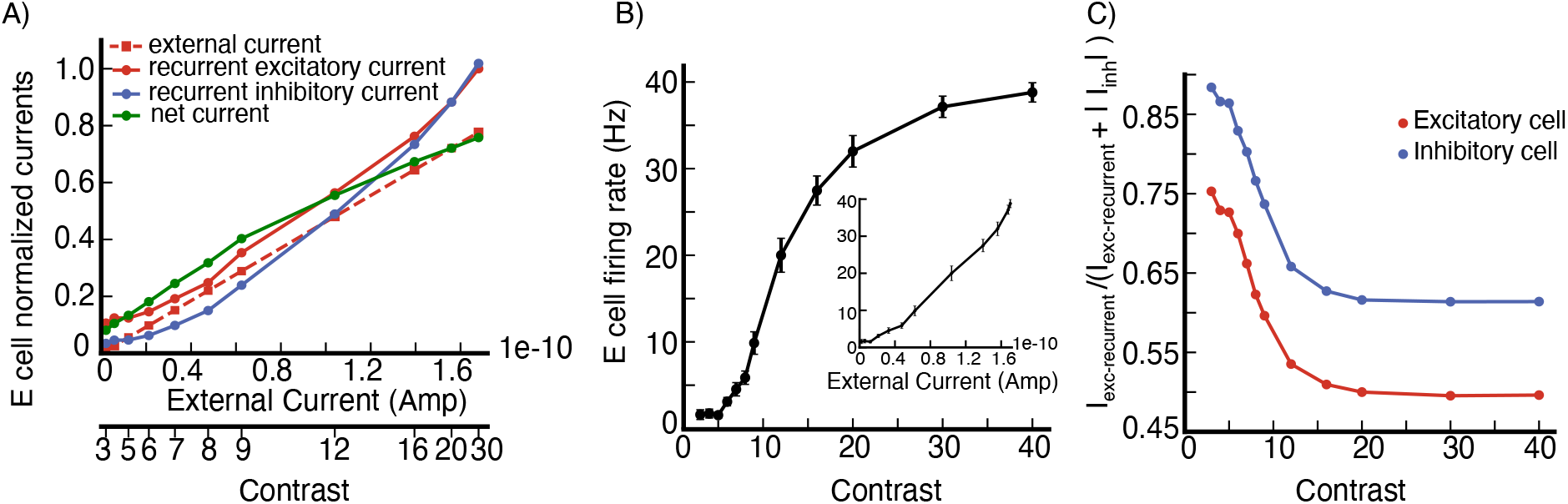
Conductance-based spiking model. (A) Input currents to an excitatory (E) cell at a randomly selected grid location (see section Model Details) vs external input current. The recurrent excitatory current *I_exc–recurrent_* = 〈*g_E_*(*R_E_* − *V*)〉_*t*_, the recurrent inhibitory current is the absolute value of *I_inh_* = 〈*g_I_*(*R_I_ − V*)〉_*t*_ and the external current *I_ext_* = 〈*g_in_*(*R_E_* − *V*)〉_*t*_ where 〈 〉_*t*_ denotes time average. The net current is (*I_ext_* + *I_exc–recurrent_* + *I_inh_*), note that *I_inh_* is negative in the spiking model. All currents are normalized to the peak value of the recurrent excitatory current. Stimulus contrast level corresponding to the external current is shown on the bottom axis. (B) Firing rate of the cell in (A) vs contrast. (C) *I_exc–recurrent_*/(*I_exc–recurrent_* + |*I_inh_*|) vs contrast for the cell in (A) and an inhibitory cell at the same grid location. In these experiments we use a stimulus with diameter 2.16° and orientation equal to the cell’s preferred orientation.

Most of our key findings in the rate model also hold in the spiking model. Both excitatory cells and inhibitory cells are surround suppressed, the excitatory cells are more strongly surround suppressed than the inhibitory cells as we see from the length tuning curves for 6 E cells (Fig. 12 top panel) and 6 I cells (Fig. 12 lower panel), and the average size tuning across 30 E cells and 30 I cells at the same grid locations (Fig. 13 A,B). The strength of surround suppression increases with increasing stimulus contrast (Fig. 12 and Fig. 13 A,B). To compute the suppressive index (SI), we fit the cells responses with a double Gaussian function. The mean SI is 0.45 ± 0.07 for the excitatory cells and 0.098 ± 0.054 for the inhibitory cells at C=10, and increases to 0.65 ± 0.09 and 0.14 ± 0.06 at C=100.

**Figure 12:**
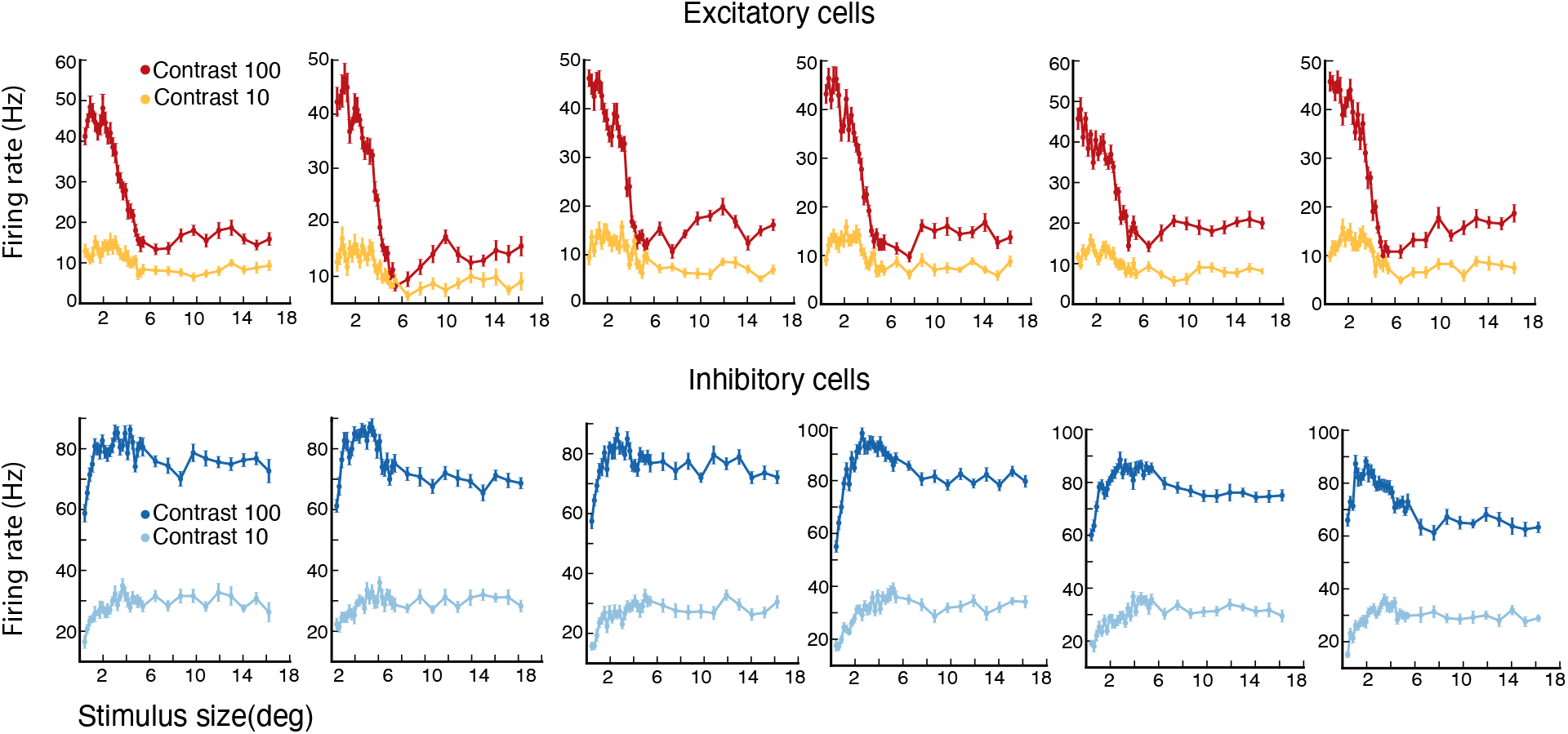
Conductance-based spiking model, surround suppression. Length tuning curves of 6 excitatory (E) cells (top panel) and 6 inhibitory (I) cells (lower panel) for two different stimulus contrast levels C=100 and C=10. The error bars are the s.e.m.

**Figure 13:**
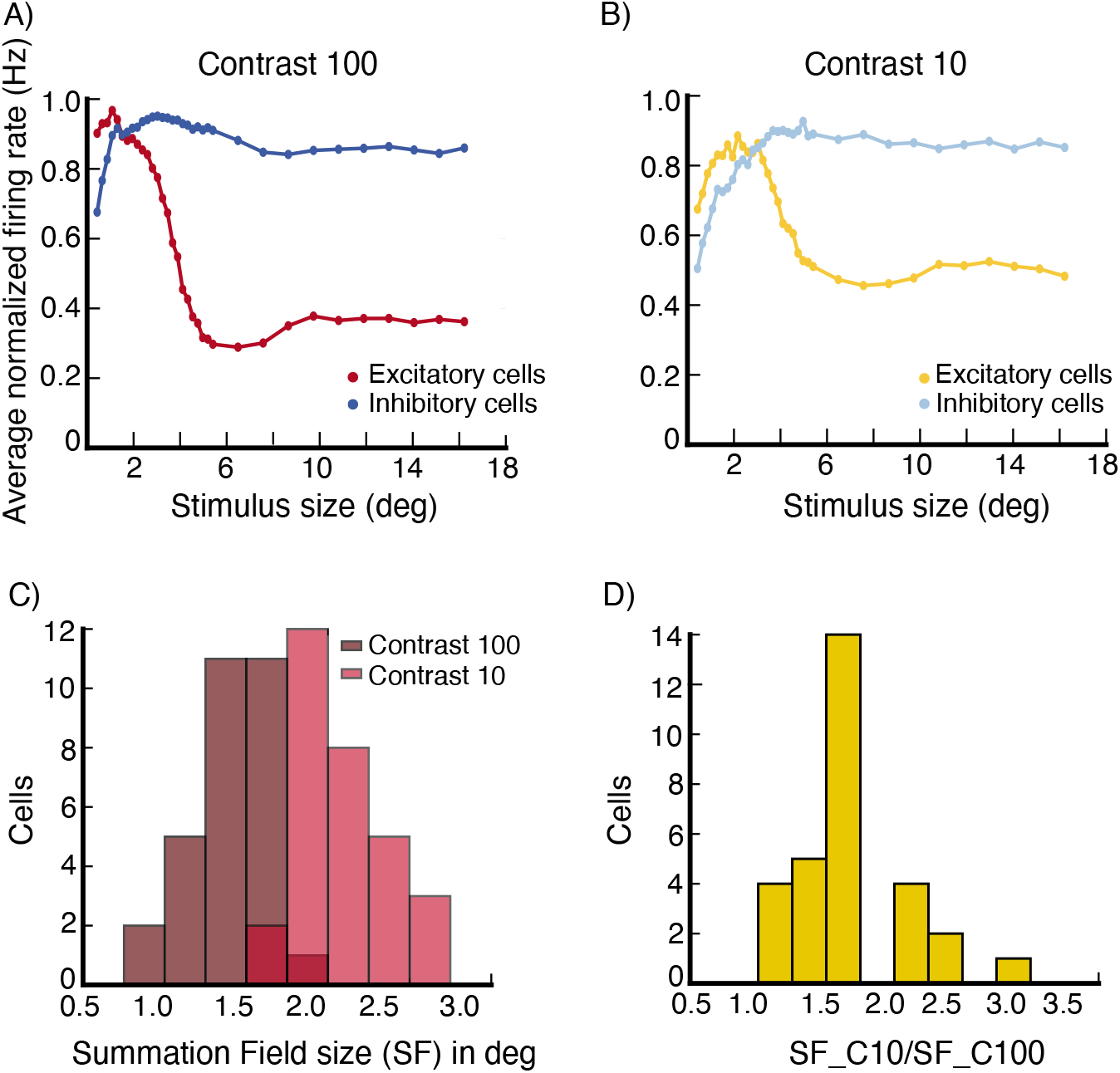
Conductance-based spiking model, surround suppression. The average firing rate of 30 excitatory (E) cells at randomly selected grid locations (see section Model Details), and of 30 inhibitory (I) cells at the same grid locations, after normalizing each cell’s rates so that its peak rate is 1.0, vs. stimulus size at contrast C=100 (A) and contrast C=10 (B). (C,D) Summation Field Sizes. (C) The distribution of summation field size of the 30 E cells used to produce panels (A) and (B) at contrasts C=100 (dark red color) and C=10 (light red color). (D) The distribution of the ratio of the summation field sizes in (C). The summation field size of all cells, is smaller at the higher contrast stimulus.

We also show the distribution of the summation field sizes of the 30 E cells selected above for two contrast levels C=100 and C=10 in Fig. 13C. The summation field size decreases with increasing contrast (Fig. 13D).

To test wether surround suppression in the spiking network is also accompanied by a decrease in excitation and inhibition, as reported by Ozeki et al. (2009), we plot the excitatory conductance values (Fig. 14A) and the inhibitory conductance values (Fig. 14B) for the same 30 E cells for a large stimulus for which all the cells are suppressed against their values for a small stimulus around which the cells respond maximally (we pick the size of the small stimulus using the same method described in Result 1). Both excitatory and inhibitory conductances are smaller for the large suppressive stimulus.

**Figure 14:**
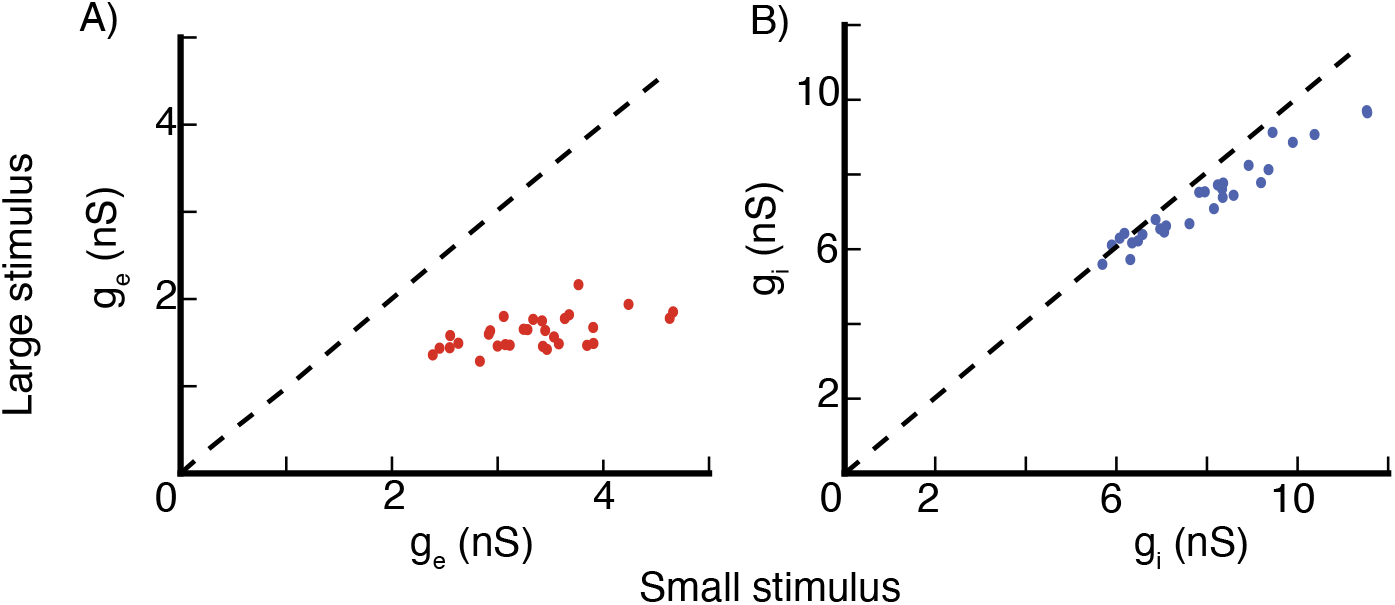
Conductance-based spiking model, surround suppression. Excitatory and in-hibitory conductances values of the 30 excitatory (E) cells used in Fig. 13. (A) Excitatory conduc-tance values of the E cells for a large suppressive stimulus are plotted against their values for a small stimulus size around which the cells respond maximally. (B) same as (A) but for inhibitory conductances. Stimulus contrast C=100.

The results for feature-specific surround suppression are qualitatively similar to what we observe in the rate model, Fig. 15B is the same as Fig. 9A,B. For surround tuning to the center orientation, even though we can see a trend in some cells similar to that we observe in the rate model, overall the phenomenon is weak for the center stimulus orientations that give a response above the spontaneous activity level, Fig. 15A shows the surround modulation map in the spiking model. We point out that we do not optimize the model parameters, so a different set of values of the connectivity profile parameters, such as the width of connectivity in orientation space and the length of connections, may lead to better results as we have observed in a few simulations. Also, the conductances values are not fine tuned, so a different set of values can for example give larger SI indexes while still producing result 3.

**Figure 15:**
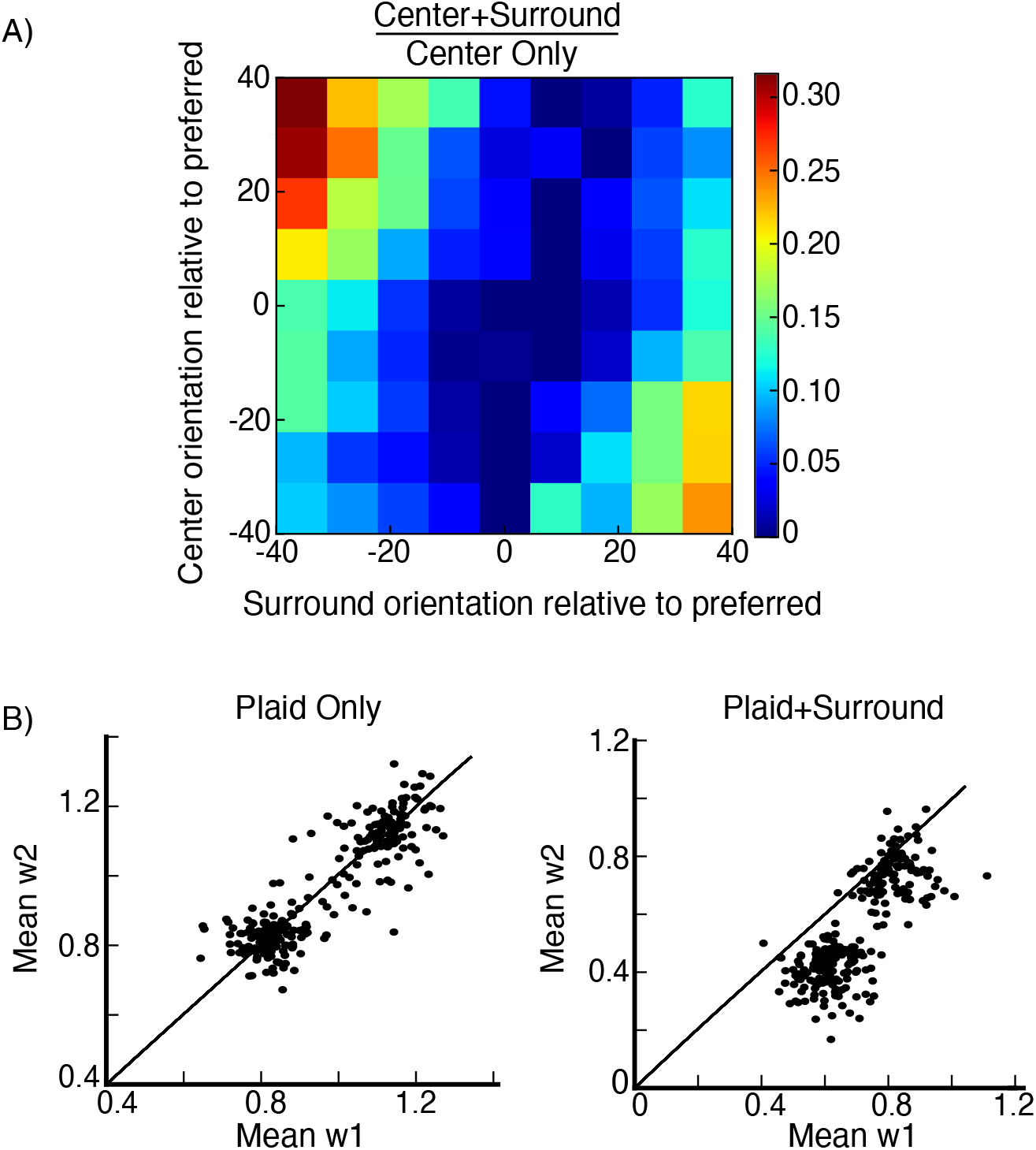
Conductance-based spiking model, surround tuning to the center orientation and feature-specific surround suppression. (A) Surround tuning to the center orientation, average surround modulation map, the data is from 23 excitatory (E) cells at randomly selected gird locations (see section Model Details), same plot as Fig. 7A. (B) Feature-specific surround suppression, the data is from 56 populations centered around 56 randomly selected grid locations (see section Model Details), same plot as Fig. 9A,B.

## Discussion

In previous work (Ahmadian et al., 2013; Rubin et al., 2015) we showed that the stabilized supralinear network motif (SSN) can explain normalization and surround suppression if combined with simple connectivity profiles, in which connection strength decreases with increasing distance across cortex or between preferred features. In Rubin et al. (2015) we presented a 2-d SSN model of V1 as a proof of principle and showed that the model can generate surround suppression. Ozeki et al. (2009) found that an iso-oriented surround stimulus reduces the values of both excitatory and inhibitory conductances of surround suppressed cells in cat V1. We have found that, unlike in the simpler models studied in Rubin et al. (2015), the 2-d model did not show this phenomena.

In this paper we built a rate-based model of layer 2/3 of V1 of animals with orientation maps and showed that lateral connections are capable of generating consistently a set of phenomena that have been observed in V1, including surround suppression, surround tuning to the center orientation and feature-specific suppression. We also showed that surround suppression is accompanied by a decrease in the excitatory and inhibitory inputs. As far as we know, this is the only spatially extended model of V1 that has shown this phenomena. The activity decay time constant for the excitatory cells in the model is fast, about the same as the single-cell time constant, as in Reinhold et al. (2015). Finally, we showed that our key results hold in a conductance-based spiking network.

The model gives insight into the circuit mechanisms that may underly the above observed phenomena. The network is a stabilized supralinear network, it has specific connectivity high-lighted by dense, strong local excitatory connections that are broadly tuned in orientation space, and long range patchy excitatory connections that are biased toward inhibitory cells at longer distances. The network is in a strongly nonlinear regime quantified by Ω*E* < 0 < Ω*i* (Ahmadian et al., 2013). The requirement that the local connectivity be broadly tuned in orientation space is essential to obtain surround tuning to the center orientation, but not to obtain surround suppression and feature-specific suppression. We note that the parameters we used, such as connectivity profile, input profile, and the underlying orientation map, were chosen without tuning them to any of the phenomena we study.

## Acknowledgments

We acknowledge computing resources from Columbia University’s Shared Research Computing Facility project, which is supported by NIH Research Facility Improvement Grant 1G20RR030893-01, and associated funds from the New York State Empire State Development, Division of Science Technology and Innovation (NYSTAR) Contract C090171, both awarded April 15, 2010. This project was supported by NIH grant R01-EY11001, grant 2016-4 from the Swartz Foundation, the Gatsby Charitable Foundation, and NSF NeuroNex Award DBI-1707398. D. Obeid would like to thank Richard Born, Alexander Trott, S. Shushruth and Cengiz Pehlevan for helpful discussions.

